# Tyrosination of α-Tubulin Controls The Initiation of Processive Dynein-Dynactin Motility

**DOI:** 10.1101/027631

**Authors:** Richard J. McKenney, Walter Huynh, Ronald D. Vale, Minhaj Sirajuddin

## Abstract

Post-translational modifications (PTMs) of α/β-tubulin are believed to regulate interactions with microtubule binding proteins. A well-characterized PTM involves the removal and re-ligation of the C-terminal tyrosine on α-tubulin, but the purpose of this tyrosination-detyrosination cycle remains elusive. Here, we examined the processive motility of mammalian dynein complexed with dynactin and BicD2 (DDB) on tyrosinated versus detyrosinated microtubules. Motility was decreased ~4-fold on detyrosinated microtubules, constituting the largest effect of a tubulin PTM on motor function observed to date. This preference is mediated by dynactin’s microtubule binding p150 subunit rather than dynein itself. Interestingly, on chimeric microtubules, DDB molecules that initiated movement on tyrosinated tubulin continued moving into a region of detyrosinated tubulin. This result indicates that the α-tubulin tyrosine facilitates initial motor-tubulin encounters, but is not needed for subsequent motility. Our results reveal a strong effect of the C-terminal α-tubulin tyrosine on dynein-dynactin motility and suggest that the tubulin tyrosination cycle could modulate the initiation of dynein-driven motility in cells.

## INTRODUCTION

Cytoplasmic dynein is the predominant minus-end directed microtubule motor in living cells (Allan, 2011; Roberts et al., 2013; Vallee et al., 2012). In metazoans, dynein is not strongly processive on its own, and long-range directional movement requires dynein’s association with the large dynactin complex, which is mediated by one of a number of coiled-coil cargo-specific adapter proteins (McKenney et al., 2014; Schlager et al., 2014). Recent structural studies suggest that the docking of dynein’s tail domain into dynactin’s actin-related 1 (Arp1) filament may reorient the dynein motor domains for productive motility (Urnavicius et al., 2015). Additionally, dynactin contains its own microtubule binding domain, located at the N-terminus of the p150^Glued^ subunit (hereafter termed p150) (Schroer, 2004), but the role of this domain in dynein motility remains unclear (Ayloo et al., 2014; Culver-Hanlon et al., 2006; Dixit et al., 2008; Kardon et al., 2009; Kim et al., 2007; Tripathy et al., 2014).

In addition to activation via an allosteric mechanism, dynein’s motor activity may also be regulated by the microtubule track. The unstructured C-terminal tail domains (CTTs) of both α- and β-tubulin are subjected to several types of PTMs, including the addition and removal of branched chains of glutamate and glycine residues, and the cyclical removal and addition of the terminal tyrosine on α-tubulin. These PTMs and genetic tubulin isotypes have been proposed to create a “tubulin-code” that could modulate interactions with molecular motors or other microtubule binding proteins (Garnham and Roll-Mecak, 2012; Janke, 2014; Janke and Bulinski, 2011; Roll-Mecak, 2015; Yu et al., 2015). In support of this idea, our previous work revealed certain selective differences in processivity and/or velocity of several kinesin motors on microtubules composed of different CTT modifications. However, the movement of yeast cytoplasmic dynein (which does not require dynactin for processive motility) revealed little selective preference for any of the microtubule substrates tested (Sirajuddin et al., 2014).

Here, we utilize in vitro reconstitution to dissect the role of the CTT domains and tubulin PTMs on the motility of mammalian dynein complexed with dynactin and the adaptor protein BicD2 (DDB). We show the α-tubulin CTT and in particular its C-terminal tyrosine are important for DDB motility.

### RESULTS

### DDB motility requires the CTT on α-tubulin but not β-tubulin

We previously reported that microtubules (MTs) treated with subtilisin, which removes the CTTs of both α- and β-tubulin, are poor substrates for single molecule DDB motility in vitro (McKenney et al., 2014). To parse the roles of the individual α- and β-tubulin CTT domains in DDB motility, we utilized a recombinant expression system in which human CTTs were fused onto the structural core of *S. cerevisiae* tubulin; the recombinant tubulin was purified by Ni-NTA affinity chromatography with a hexahistidine tag placed within a loop on α-tubulin that faces the MT lumen (Sirajuddin et al., 2014). For these assays, we used the CTTs corresponding to human tubulin isoforms TUBA1A and TUBB2A, which are common isoforms expressed in mammals. The DDB complex was purified from porcine brain; a SNAPf tag was fused to the truncated recombinant BicD2 so that the movement of the DDB complex could be followed by total internal reflection fluorescence (TIRF) microscopy (McKenney et al., 2014). Single molecules of DDB moved processively on microtubules polymerized from recombinant TUBA1A/TUBB2A tubulin (hereafter referred to as WT tubulin) with a velocity similar to that reported previously on mammalian MTs (Fig. 1A-B)(McKenney et al., 2014; Schlager et al., 2014).

**Figure 1.**
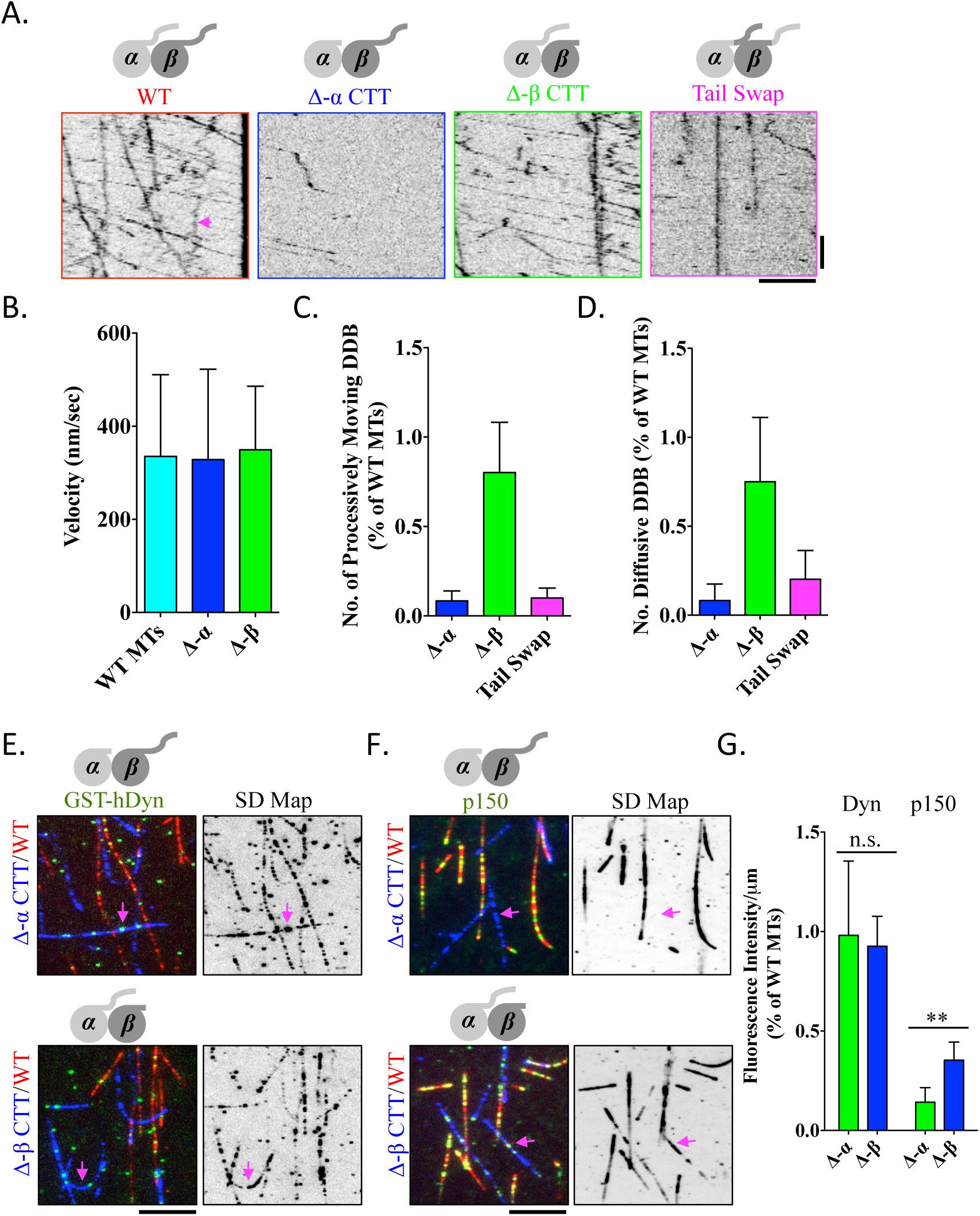
Role of tubulin tail domains in DDB movement. **(A)** Representative kymographs showing tetramethylrhodamine (TMR)-labeled DDB movement on each type of recombinant MT lattice, as indicated. Scale bars, 5 μm and 25 sec. Arrowhead marks a diffusive DDB complex. See also Video 1. **(B)** Quantification of DDB velocity on MT lattices lacking each of the tubulin-CTTs (n ≥ 75 DDB complexes per condition from at least two independent experiments). **(C-D)** Quantification of the number of processive **(C)**, or diffusive **(D)** DDB complexes relative to WT MTs in the same chamber. n ≥ 30 MTs quantified for each condition from at least two independent experiments. Processive (unidirectional motion exceeding 0.5 μm) and diffusive (back- and-forth motion) events were scored in a field of view and then divided by the lengths of MTs and time, thus normalizing to a value of number of motility events per μm MT per sec (n ≥ 30 MTs quantified for each condition from at least two replicate experiments). Error bars SD. **(E-F)** TIRF images of 1nM TMR-labeled GST-hDyn **(E)**, or 0.3 nM TMR-labeled p150 **(F)** molecules (green) bound to either Δα-CTT (top), or Δβ-CTT MTs (bottom). Right, standard deviation maps from an entire time sequence reveal ensemble binding and dissociation events that lead to variations in pixel intensities (see Methods). Scale bars are 5 μm. **(G)** Quantification of mean fluorescence intensity (arbitrary units) per μm MT for GST-hDyn or p150 bound to the indicated mutant MT relative to WT MTs in the same chamber. Image shows first frame from time series. The intensity of GST-hDyn was quantified from maximum intensity projections of the entire time sequence due to transient binding of GST-hDyn to the MT (n ≥ 40 MTs quantified for each condition from at least two replicate experiments). Error bars represent SD. P value (unpaired *t* test) = 0.0037 for p150.

Next, we prepared recombinant tubulin lacking either the α- or β-tubulin CTT. DDB motility was assayed in chambers containing the Δ-CTT MTs as well as control WT MTs (each labeled with ~1% incorporation of fluorescently-labeled and biotinylated porcine brain tubulin). We observed a dramatic decrease (~85%) in the number of processive DDB molecules on MTs lacking the α-CTT compared to WT MTs in the same chamber (Fig. 1A, C, Video S1). Strikingly, the frequency of processive DDB movement on MTs lacking the β-CTT was only modestly reduced (~20%) compared to WT MTs (Fig. 1A, C, Video S1). While deletion of the α-CTTs strongly suppressed DDB movement, the few complexes that did move on these MTs did so at a similar velocity as WT MTs (Fig. 1B). As previously reported, a subset of DDB complexes (~30%) displayed only diffusive back-and-forth motion on the MT with no obvious directional bias (Fig. 1A, arrow, (McKenney et al., 2014). Similar to processive complexes, the number of diffusive DDB complexes was dramatically reduced by deletion of the α-CTT domain, but not the β-CTT domain (Fig. 1A, D). Thus, the α-CTT, but not the β-CTT domain, plays a major role in both processive and diffusive DDB interactions with the MT lattice.

We next swapped the positions of the CTTs, such that the α-tubulin CTT was placed on β-tubulin and vice versa (see Methods). Strikingly, both processive and diffusive DDB movement was dramatically reduced on these MTs, despite the presence of the α-CTT on the β-tubulin core (Fig. 1A, C-D, Video S1). These results demonstrate that the DDB interaction with the MT lattice requires the stereospecific location of the α-tubulin CTT.

### Analysis of the MT-binding domains of dynein and dynactin

DDB contains two defined MT-binding domains; one located within the dynein motor domain, and the other on the p150 subunit of dynactin. We investigated how each of these MT-binding domains interacts with the CTTs. We tested the previously characterized recombinant GST-dimerized human dynein motor domain (GST-hdyn) and a truncated dimeric construct of the neuronal isoform of p150 that contains the CAP-Gly and basic domains, both of which have been shown to interact with MTs (Ayloo et al., 2014; Culver-Hanlon et al., 2006; McKenney et al., 2014; Tripathy et al., 2014; Trokter et al., 2012).

In the presence of ATP, GST-hDyn bound transiently, but did not move processively on our recombinant MTs, similar to previous reports on mammalian MTs (McKenney et al., 2014; Torisawa et al., 2014; Trokter et al., 2012). A map of the standard deviation of the fluorescence intensity for each pixel over an entire time series was generated to provide insight into the ensemble binding events of GST-hDyn (Cai et al., 2009; Cai et al., 2007). This analysis revealed that motor binding was not perturbed by the deletion of either the α- or β-tubulin CTT (Fig. 1E). Quantification of GST-hDyn fluorescence on the MT also revealed that motor binding was not significantly diminished by the loss of either CTT (Fig. 1G). These results are consistent with structural studies suggesting that dynein binds to the MT lattice at the interface of the α/β tubulin dimer and does not make contact with the tubulin CTTs (Carter et al., 2008; Mizuno et al., 2004; Redwine et al., 2012; Uchimura et al., 2015). While mammalian dynein’s non-processive interaction with the MT was not affected by loss of the tubulin CTTs in our assay, previous studies reported that removal of the CTTs leads to a decrease in processive run-lengths for yeast dynein (Redwine et al., 2012; Sirajuddin et al., 2014), as well as for mammalian dynein attached to plastic beads (Wang and Sheetz, 2000).

The p150 construct also bound to the recombinant MT lattice and displayed bidirectional and diffusive motility as previously described (Ayloo et al., 2014; Culver-Hanlon et al., 2006; Tripathy et al., 2014). In contrast to GST-hDyn, p150 binding was strongly diminished by removal of the α-CTT, but less so by removal of the β-CTT (Fig. 1F-G). Thus the p150 MT-binding domain, similar to DDB, displays a strong requirement for the presence of the α-CTT, while the dynein-MT interaction is not affected by the absence of either CTT domain.

### The C-terminal tyrosine of α-tubulin is critical for DDB motility

Because of the strong requirement for the α-CTT in DDB motility, we turned our attention to PTMs specific to this domain. One of the first and most prominent PTMs to be discovered is the cyclical removal and re-addition of the terminal tyrosine residue on the α-CTT (Arce et al., 1975; Gundersen et al., 1984; Hallak et al., 1977). Interestingly, the N-terminal CAP-Gly domain of p150 has been previously shown to recognize the – EEY/F motif at the C-terminus of α-tubulin (Peris et al., 2006; Wang et al., 2014). Thus, we wanted to test whether this tyrosine residue plays an important role in the DDB-MT interaction.

We assayed DDB motility on recombinant tubulin in which we genetically deleted the terminal tyrosine residue on the α-CTT (detyrosinated MTs). We note that tubulin purified from yeast does not contain other types of PTMs that are commonly found in tubulin purified from mammalian sources (polyglutamylation, polyglycylation, acetylation), and budding yeast lack the ligase that is needed to post-translationally add tyrosine to the C-terminus of α-tubulin. Strikingly, DDB movement was ~4-fold more frequent on MTs containing tyrosinated α-CTT than on detyrosinated MTs placed in the same chamber (Fig. 2A-B, Video S2). Maps of the pixel standard deviation over the whole acquisition series confirmed that DDB bound much less frequently on detyrosinated MTs than on tyrosinated MTs (Fig. 2A). Similarly, the diffusive population of DDB was strongly reduced on detyrosinated MTs compared to tyrosinated MTs (Fig. 2C). We next polymerized MTs containing variable amounts of tyrosinated α-tubulin and quantified the frequency of DDB movement (see Methods). This analysis revealed that maximal DDB motility is achieved when 50% of the MT lattice contains tyrosinated α-tubulin, while ~75% of full motility is achieved at 30% tyrosination (Fig. 2D). In summary, these results demonstrate that the C-terminal tyrosine on α-tubulin is necessary for a robust DDB-MT interaction.

**Figure 2.**
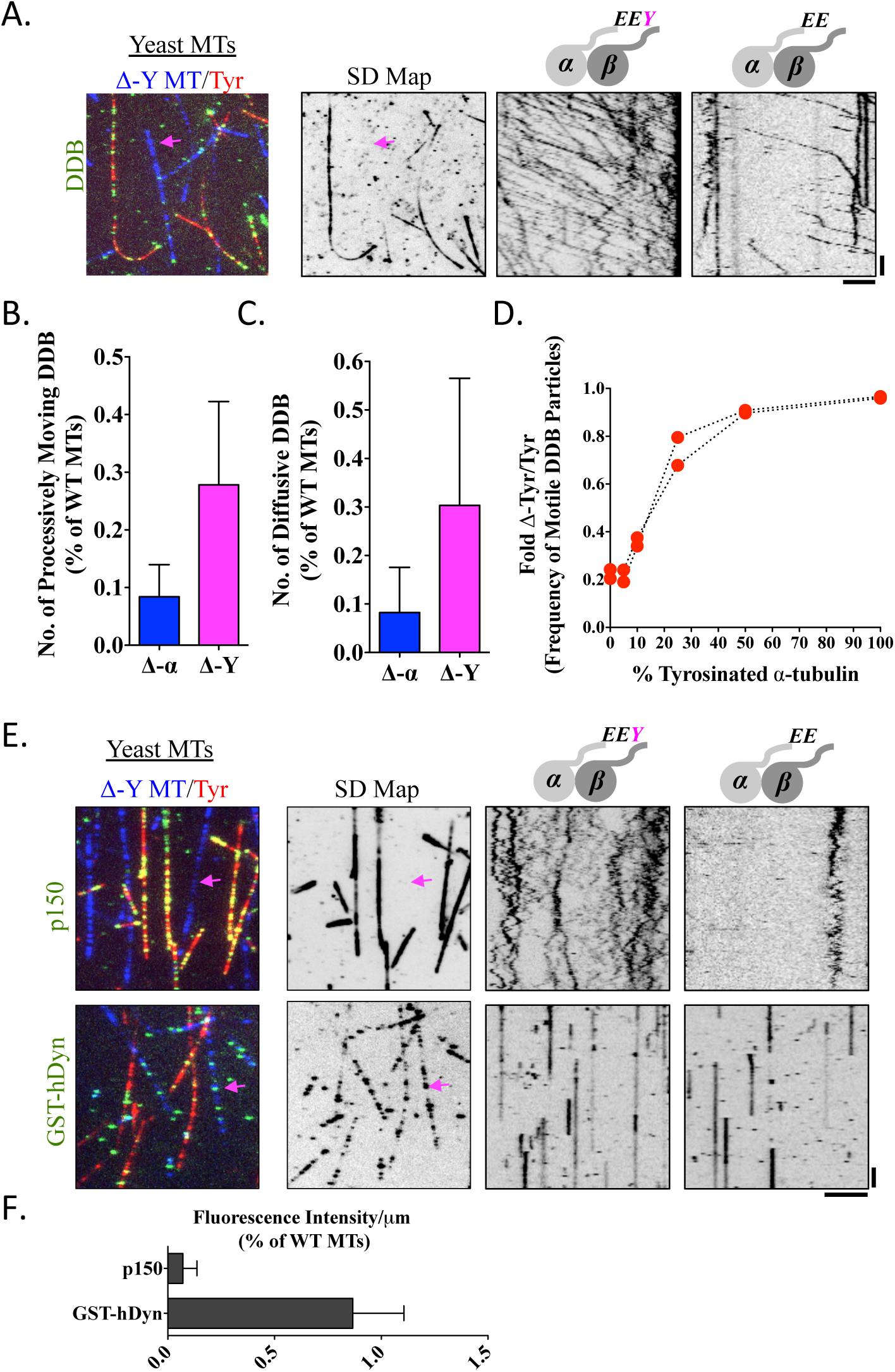
Requirement of α-tubulin tyrosination for processive DDB movement. **(A)** TIRF image of TMR-labeled DDB molecules bound to either tyrosinated (red) or detyrosinated (blue) MTs. A standard deviation map of the whole time sequence is shown, revealing the preferential association of DDB with tyrosinated MTs versus detyrosinated MTs (arrow). Representative kymographs of DDB movement are shown from each type of MT present in the same chamber. Scale bars are 5 μm and 25 sec. See also Video 2. **(B-C)** Quantification of the number of processive **(B)** or diffusive **(C)** DDB complexes per μm MT per sec on detyrosinated MTs relative to tyrosinated MTs in the same chamber (n ≥ 30 MTs quantified for each condition). Error bars represent SD. Data from Δα-CTT MTs (Fig. 1) are re-plotted for comparison. **(D)** Quantification of the frequency of processive DDB molecules as a function of the amount of tyrosinated tubulin incorporated into the lattice normalized to fully tyrosinated MTs. Data points from two independent experiments are shown. **(E)** TIRF images of 0.3 nM TMR-labeled p150 (top) or 1 nM TMR-labeled GST-hDyn (bottom) molecules bound to MTs either Δα-CTT (top), or Δβ-CTT (bottom). Right, standard deviation maps of protein binding over an entire time sequence and kymographs from either tyrosinated or detyrosinated are shown. Scale bars are 5 μm and 25 sec. **(F)** Quantification of mean fluorescence intensity (arbitrary units) per μm MT for TMR-labeled GST-hDyn or p150 bound to the indicated mutant MT relative to WT MTs in the same chamber. The intensity of GST-hDyn was quantified from maximum intensity projections of the entire time sequence due to transient binding of GST-hDyn to the MT. (n ≥ 40 MTs quantified for each condition from at least two independent experiments). Error bars represent SD.

We next examined how the C-terminal α-tubulin tyrosine affects the dynein- and p150-MT interactions individually. GST-hDyn showed no preference for binding to tyrosinated- or detyrosinated MTs (Fig. 2E-F), which is consistent with the above findings (Fig. 1E-G), and previous reports for mammalian and yeast dynein (Carter et al., 2008; Redwine et al., 2012; Sirajuddin et al., 2014; Uchimura et al., 2015). In striking contrast, removal of the terminal tyrosine nearly abolished the interaction of p150 with the detyrosinated MT (Fig. 2E-F). We conclude that in the absence of other tubulin PTMs, tyrosination of the α-CTT is critical for both DDB motility and p150 binding, but not for dynein binding. These results suggest that the MT binding requirements of p150 dictate DDB movement along the MT lattice.

To probe the role of other tubulin isotypes and PTMs in conjunction with the α-CTT tyrosination state in DDB motility, we turned to tubulin purified from porcine brain, which is a composite of many tubulin isotypes and PTMs. Treatment with the enzyme carboxypeptidase A (CPA) specifically removed the α-CTT tyrosine without affecting other PTMs (Fig. 3A) (Webster et al., 1987b). Strikingly, CPA treatment reduced the number of processively moving DDB molecules on digested MTs by ~50% compared to WT MTs in the same chamber (Fig. 3B-C). Examination of both GST-hDyn and p150 binding to CPA MTs showed CPA treatment had little effect on the binding of GST-hDyn, but strongly reduced the binding of p150 to the MT (Fig. 3D-E), likely due to the loss of interaction between p150’s CAP-Gly domain and the α-CTT –EEY/F motif. Thus, removal of the α-CTT tyrosine, in the background of mixed tubulin isotypes and PTMs, reduces both DDB movement and p150 binding.

**Figure 3.**
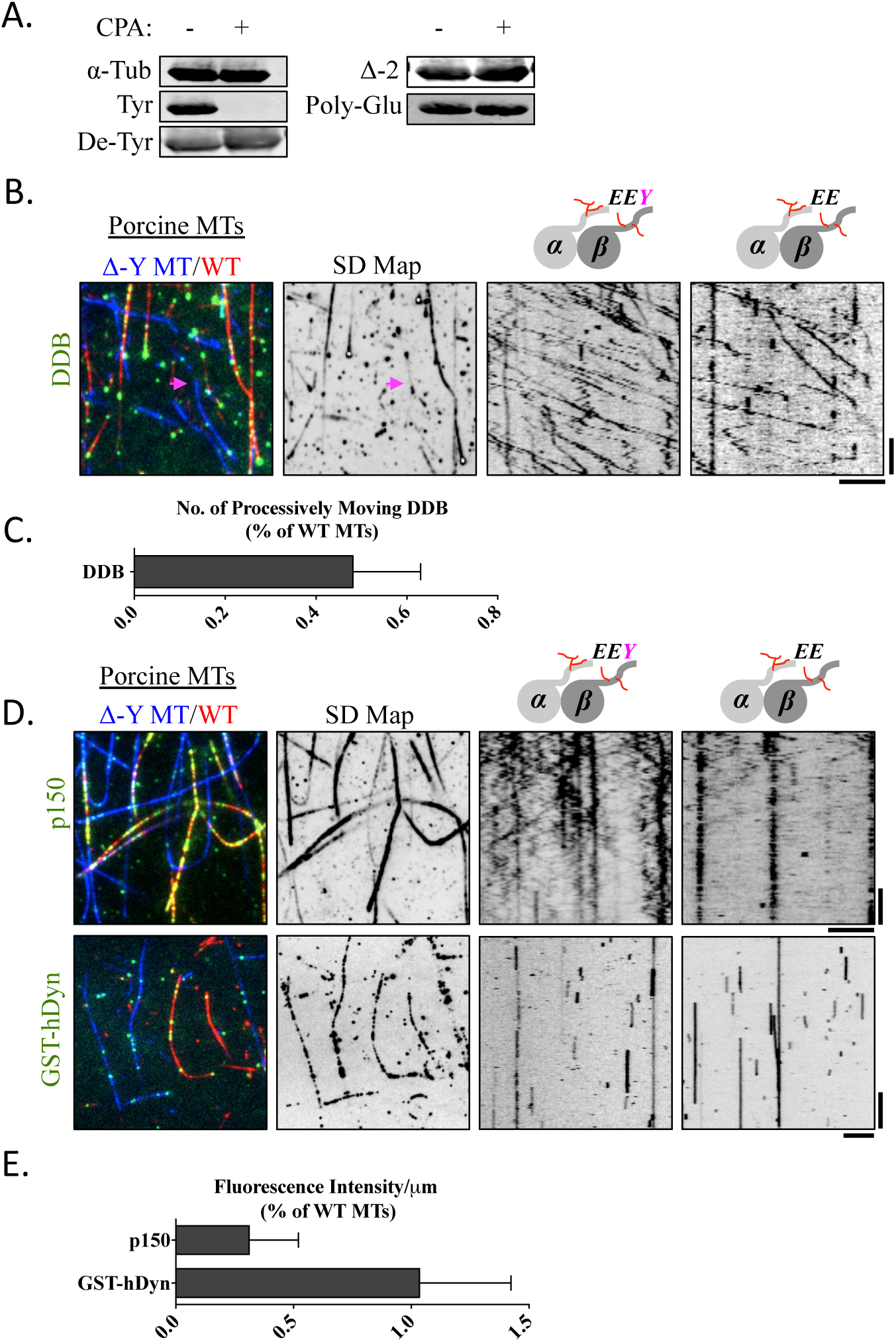
Role of tyrosination in DDB movement on brain MTs. **(A)** Immunoblot of brain tubulin with our without incubation with carboxypeptidase A (CPA). Note CPA specifically removes the α-tubulin C-terminal tyrosine without affecting other PTMs. **(B)** TIRF image of TMR-labeled DDB molecules (green) on either WT (red) or CPA-treated (blue) pig brain MTs. Standard deviation map of the entire time sequence shows DDB preference for WT MTs and kymographs from either WT or CPA-treated are shown. Scale bars are 5 μm and 25 sec. **(C)** Quantification of the number of processive DDB complexes per μm MT per sec. on the CPA-treated MTs, relative to WT MTs in the same chamber (n ≥ 100 of each type of MT). Error bars represent SD. **(D)** TIRF images of 0.3nM TMR-labeled p150 (top) or 1nM TMR-labeled GST-hDyn (bottom) molecules bound to either WT (red), or CPA-treated MTs (blue). Right, standard deviation maps of protein binding over an entire time sequence and kymographs from either WT or CPA-treated are shown. Scale bars are 5 μm and 25 sec for p150, and Scale bars, 5 μm and 12 sec for GST-hDyn **(E)** Quantification of mean fluorescence intensity (arbitrary units) per μm MT for TMR-labeled GST-hDyn or p150 bound to CPA-treated MTs relative to WT MTs in the same chamber. The intensity of GST-hDyn was quantified from maximum intensity projections of the entire time sequence due to transient binding of GST-hDyn to the MT (n ≥ 40 MTs quantified for each condition from at least two replicate experiments). Error bars represent SD.

To examine whether the p150 CAP-Gly is directly required for p150’s preference for tyrosinated MTs, we purified a neuron-specific isoform of p150 that lacks the N-terminal CAP-Gly domain (termed p135), but retains a portion of the adjacent basic domain that also bind MTs (Fig. S1A)(Culver-Hanlon et al., 2006; Tokito et al., 1996). At similar protein concentrations, much less p135 protein bound along MTs compared to p150, consistent with its previously reported lower affinity for MTs (Fig. S1B)(Lazarus et al., 2013)). In contrast to p150, p135 did not bind preferentially to tyrosinated MTs (Fig. S1B-C), revealing that the CAP-Gly domain dictates p150’s preferential interaction with tyrosinated MTs. Our results suggest that recognition of the α-tubulin tyrosine by the CAP-Gly domain of p150 plays a key role for DDB motility, even in the background of a heterogeneous mixture of tubulin isotypes and other PTMs.

### Initiation, but not the continuation of processive DDB motility requires α-tubulin tyrosination

Having established that tyrosination of the α-CTT is critical for both p150 binding and DDB movement, we next sought to investigate how p150 may regulate DDB motility. Dynactin has been proposed to tether dynein to the MT surface during processive motility (Ayloo et al., 2014; Culver-Hanlon et al., 2006; King and Schroer, 2000; Ross et al., 2006). We reasoned that if this model were correct, we should observe a cessation of processive movement when a DDB complex moving processively along a tyrosinated stretch of a MT encountered a detyrosinated section of a MT. To create such a bipartite MT lattice, stabilized MTs composed of either tyrosinated or detyrosinated α-tubulin were joined end-on-end through a spontaneous annealing reaction (Rothwell et al., 1986). This resulted in single chimeric MTs containing localized zones of tyrosinated and detyrosinated tubulin (Fig. 4A). We then observed the behavior of processive DDB complexes as they traversed a junction between a tyrosinated and detyrosinated MT segment. DDB molecules usually initiated movement on tyrosinated regions of a chimeric microtubule, as described earlier (Fig. 2). Surprisingly, the majority of processive DDB complexes moved uninterrupted through the tyrosinated to detyrosinated MT boundary and continued long processive movements along the detyrosinated section of the MT (59.2%, Fig. 4B, Video S3), while the remaining DDB complexes either stalled at the junction or dissociated from the MT. In contrast to this behavior, the subset of diffusive DDB complexes never traversed the intersection, and often accumulated there (Fig. 4B, arrow). Similarly, the p150 construct bound and diffused nearly exclusively along the tyrosinated section of MT and rarely crossed the boundary to a detyrosinated section of MT (Fig. 4C). Because the diffusive population of DDB complexes behaved similarly to p150 with respect to the tyrosinated boundary, we speculate that the diffusive DDB complexes are bound to the MT exclusively through the p150 subunit of dynactin.

**Figure 4.**
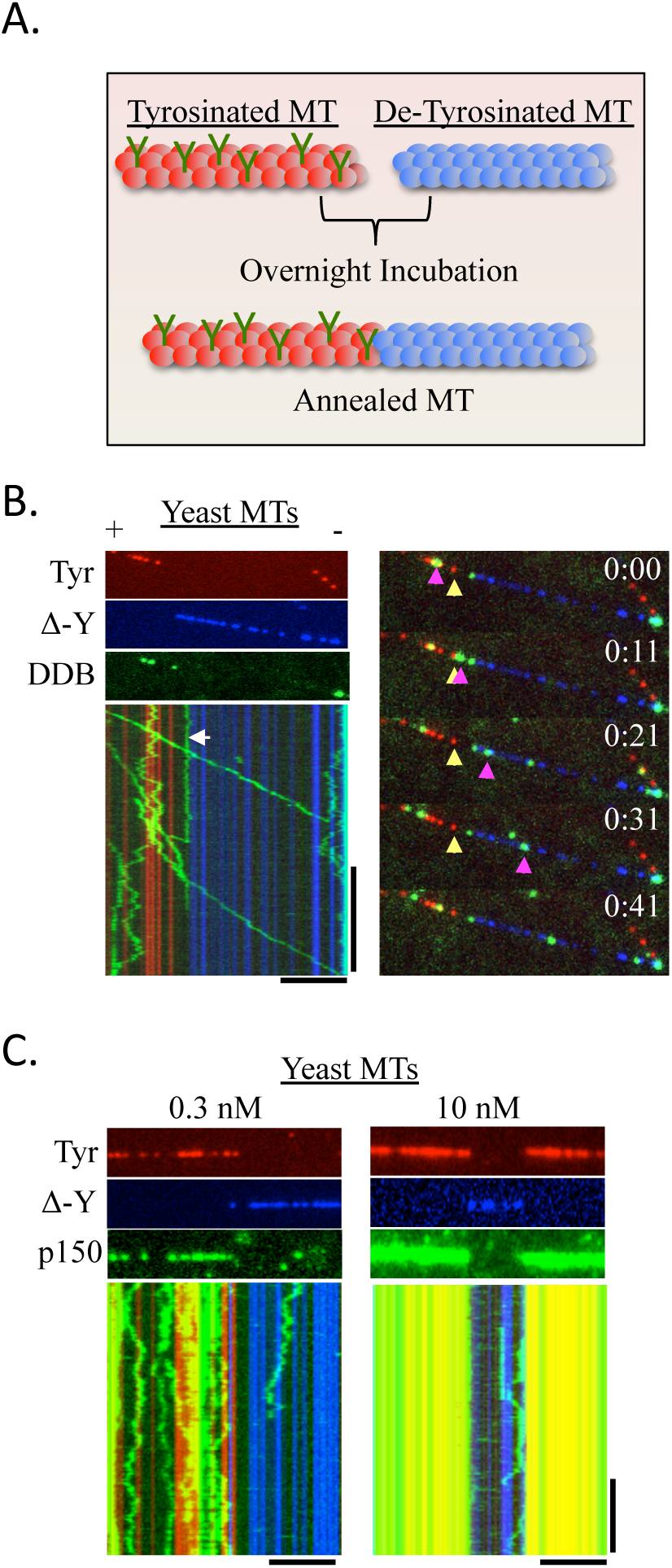
α-Tubulin tyrosination is required for the initiation, but not continuation of processive DDB movement. **(A)** Schematic showing the assembly of a chimeric MT lattice created by end-to-end annealing of stabilized MTs. The MTs are differentially fluorescently labeled. Scale bars are 5 μm and 25 sec. See also Video 3. **(B)** Example of a chimeric MT generated from recombinant yeast tubulin containing labeled tyrosinated (red) and detyrosinated (blue) sections. TMR-labeled DDB molecules (green) move processively through the annealed junction between the two types of MTs (59.2%, n=574 processive complexes, 51 MT-MT junctions, 2 independent experiments). Diffusive DDB molecules (white arrow) are primarily observed on the tyrosinated section of the MT and do not diffuse across the boundary. Right, movie frames showing a processive DDB complex (pink arrowhead) moving across the junction (yellow arrowhead) of tyrosinated and detryosinated MT. The (+) and (−) signs denote MT polarity inferred from the directionality of DDB movement. **(C)** TIRF images of low (0.3 nM) and high (10 nM) concentrations of TMR-labeled p150 (green) bound to chimeric yeast MTs containing tyrosinated (red) and detyrosinated (blue) sections of MT. Representative kymographs below show diffusive behavior of p150 molecules on the MT. Note that p150 rarely diffuses into the detyrosinated section of MT, even at high concentrations. All scale bars are 5 μm and 10 sec.

We further confirmed these results using annealed MT lattices composed of sections of porcine tubulin differentially treated with CPA to remove the α-CTT tyrosine. First, processive DDB complexes traversed the annealed boundary between two WT MTs without pausing (93.5%, Fig. S2A). Next, we observed that most processive DDB complexes were able to traverse the boundary from WT tubulin into a section of CPA-treated tubulin (77.6%, Fig. S2B). Finally, we created a single annealed MT composed of untreated and subtilisin digested porcine tubulin, which removes both CTTs (Δ-CTT) and largely abolished DDB motility (McKenney et al., 2014). Our results show that a subset of processive DDB complexes that bound initially to the untreated tubulin section of the MT crossed the boundary and continued processive motility on the Δ-CTT section of MT (41.1%, Fig. S2C). As p150 cannot interact with Δ-CTT MTs (Lazarus et al., 2013; McKenney et al., 2014), we reason that p150 does not act as a continuous MT tether during processive movement of DDBs on Δ-CTT MTs. From these data, we conclude that the initiation of processive DDB motility, but not its continuation, requires the interaction of p150 with the C-terminal tyrosine of α-CTT.

## DISCUSSION

We have found that the initiation of processive motility by dynein-dynactin is enhanced by the interaction of the CAP-Gly domain of p150 with the C-terminal tyrosine of α-tubulin; however, the subsequent continuation of processive motility does not require this interaction. These results, in conjunction with other recent findings, help to formulate a model for how dynactin might regulate motility of mammalian dynein. Previous work has indicated that p150 in the dynactin complex may not interact with MTs (Kardon et al., 2009; McKenney et al., 2014), likely due to a folded-back, autoinhibited state that places the CAP-Gly domain close to the Arp1 backbone of dynactin (Tripathy et al., 2014; Urnavicius et al., 2015). However, the interaction of dynein with an adapter protein (e.g. BicD2) may release this inhibition and allow p150 to interact with MTs (Urnavicius et al., 2015). In fact, in the DDB complex, p150 may provide the dominant means of initiating the first encounter with the MT. Evidence for this is that dynein alone interacts well with detyrosinated MTs, while the entire DDB complex shows a strong preference for tyrosinated over detyrosinated MTs. The initiation of the p150-MT encounter may provide time or steric space for both microtubule-binding domains of dynein to engage with the MTs; once this occurs, our data suggests that the p150-MT interaction is no longer required. Dynactin-BicD2, however, may serve an additional allosteric role in orienting the two motor domains of dynein homodimer to facilitate processive movement (Chowdhury et al., 2015; Urnavicius et al., 2015). These results and model also might explain the roles for two isoforms of p150 in neuronal tissues- the p150 isoform discussed above and the p135 isoform lacking the CAP-Gly domain (Tokito et al., 1996). Dynactin molecules with either isoform may be equally effective in inducing the allosteric change of the dynein homodimer, but p135-containing complexes would have a lower initiation rate of motility (McKenney et al., 2014) and lack a preference for tyrosinated MTs.

How might the preference of dynein-dynactin for tyrosinated α-tubulin influence cargo transport in vivo? In cells, many cargos bind plus-end directed kinesin and minus-end directed dynein motors simultaneously, resulting in often salutatory, bidirectional motion. Interestingly, several in vivo studies have suggested that certain kinesin motors preferentially move on detyrosinated MTs in vivo (Cai et al., 2009; Konishi and Setou, 2009). In vitro, the processivity of kinesin-2 is also enhanced ~2-fold on detyrosinated MTs (Sirajuddin et al., 2014). The preference of dynein-dynactin for tyrosinated MTs and kinesins for detyrosinated MTs could, in principle, enable the post-translational modification to bias the direction of transport. For example, a cargo with a mixture of dyneins and kinesins (Hendricks et al., 2010; Rai et al., 2013; Soppina et al., 2009) could be biased towards the periphery (MT plus end) if interacting with a heavily detyrosinated MT, or conversely toward the cell center (MT minus end) if it encounters a tyrosinated MT. Another function of this tyrosination preference might involve regulating in vivo motility through interactions of p150 with microtubule plus-end binding proteins. Previous studies have found that the p150 CAP-Gly domain also recognizes the C-terminal –EEY/F motifs on EB1 and CLIP-170 (Duellberg et al., 2014; Honnappa et al., 2006; Lansbergen et al., 2004; Weisbrich et al., 2007), which might provide an additional mechanism for facilitating the initiation of dynein motility at the plus-ends of MTs (Lloyd et al., 2012; Moughamian and Holzbaur, 2012).

In summary, our results provide strong support for the “tubulin code” hypothesis in regulating the interaction of motor proteins with MTs. Our observations of DDB preference for tyrosinated MTs constitute the largest effect of PTMs on motor activity in vitro reported to date (Barisic et al., 2015; Sirajuddin et al., 2014). An upcoming challenge will be to decipher how motor preferences for certain MT tracks translate into controling the frequency and/or directionality of cargo transport in living cells.

## METHODS

### Recombinant tubulin, polymerization and chimeric microtubules

Recombinant yeast tubulin with human C-terminal tails (chimeric yeast core-human CTT tubulin heterodimer) contains an internal hexahistidine tag located on the α-tubulin subunit and was expressed and purified as described earlier (Sirajuddin et al., 2014). The tail-swapped tubulin used here was constructed by replacing the α-CTT with β-CTT and vice versa. For the yeast alpha tubulin core fused to TUBB2A CTT, the 19 amino acid peptide DATADEQGEFEEEEGEDEA from TUBB2A was genetically fused to yeast alpha tubulin after amino acid 420. Similarly the 13 amino acid peptide SVEGEGEEEGEEY from TUB1A was genetically fused to yeast beta tubulin after amino acid 426. The purified recombinant tubulin heterodimer (~1-2 mg/ml concentration) was stored in BRB80 buffer (80 mM Pipes pH 6.8, 2 mM MgCl_2_, 1 mM EGTA) and 200 μM GTP at -80° C. For each day of assays, the recombinant tubulins were polymerized overnight at 30° C, in the presence of 2 mM GTP with 5 μM Epothilone-B (Sigma). All the polymerized microtubules contain ~1:250 and ~1:100 ratio of biotinylated and fluorescently labeled (Alexa-488 or -640) porcine brain tubulin respectively. The percentage ratios of tyrosinated microtubules reported in Figure 2D were prepared by mixing the following molar ratios of tyrosinated and detyrosinated recombinant tubulin; 0.05:1, 0.1:1, 0.5:1, 1:1 representing 5, 10, 25, 50 percent tyrosinated microtubules respectively.

Porcine brain tubulin was purified according to standard methods (Castoldi and Popov, 2003). To polymerize brain MTs, purified brain tubulin was first incubated with 1 mM GTP at 37^o^ C for 10 min, followed by the addition of 20 μM taxol for an additional 20 min. Polymerized MTs were purified further by centrifugation at 22,000 x g for 10 min over a 25% sucrose cushion made in BRB80 buffer containing 10 μM paclitaxel. The carboxypeptidase A (CPA) treated porcine brain tubulin protocol was adapted from (Webster et al., 1987a). Briefly, 12 μg/ml CPA (Sigma) was incubated with tubulin (12.5 mg/ml) and 1 mM GTP for 20 min at 37^o^ C, followed by the addition of 20 μM taxol for an additional 20 min. This was the lowest concentration of protease that completely remove the signal by western blotting with an antibody specific for tyrosinated form of tubulin. The digestion was stopped by the addition of 10 mM DTT and the CPA enzyme was removed by centrifugation of the MTs over a 25% sucrose cushion as described above. The subtilisin treated MTs were prepared as described earlier (McKenney et al., 2014). Chimeric microtubules were prepared by mixing equal parts of two species of polymerized microtubules, labeled with different fluorescent dyes, and incubated overnight at 30 ^o^ C for recombinant yeast MTs or room temp for porcine brain MTs.

### Antibodies

Antibodies used were: anti-alpha tubulin DM1A (Sigma), anti-tyrosinated tubulin (T9028, Sigma), anti-detyrosinated tubulin (ab48389, Abcam), anti-Delta2 tubulin (AB3203, Milipore), and anti-polyglutamated tubulin GT335 (Adipogen). Western blots were visualized using a LiCor Odyssey system.

### Purification of dynein-dynactin-BicD2 (DDB) complex and P150 constructs

Recombinant strepII-SNAPf-BicD2 and p150-sfGFP-SNAPf-strepII were purified from bacteria as previously described (McKenney et al., 2014). Human p135 cDNA was obtained from the mammalian gene collection (Dharmacon, GE, GenBank accession number BC071583.1). A construct encoding the first 413 amino acids of human p135, followed by an sfGFP-SNAPf-StrepII cassette was constructed similarly in a pET28 backbone (McKenney et al., 2014). Proteins were expressed from bacteria grown in LB media and induced with 1mM IPTG for 18 hours at 18^o^C. All bacterially expressed proteins were purified by affinity chromatography on Streptactin resin (GE Life Sciences) followed by gel filtration on a Superose 6 column (GE Life Science). SNAPf-BicD2 was purified and gel filtered in buffer A (30 mM Hepes pH 7.4, 50 mM K-acetate, 2 mM Mg-acetate, 1 mM EGTA, 10% glycerol), with protease inhibitor cocktail (Promega) and 2 mM DTT. Both p150 and p135 constructs were purified in Buffer B (50 mM Tris-Base pH 8.0, 150 mM K-acetate, 2 mM Mg-acetate, 1 mM EGTA, 10% glycerol), with protease inhibitor cocktail and 2 mM DTT. The purified proteins were then gel filtered on a Superose 6 column equilibrated in buffer A. Peak fractions were pooled, concentrated on Amicon filters (Millipore), and flash frozen in LiN_2_.

Recombinant SNAPf-GST-hDyn protein was prepared using the Bac-to-Bac baculovirus system (Invitrogen) as previously described (McKenney et al., 2014). The purified protein was labeled with 10 μM SNAP-Cell TMR-Star (NEB) while bound to the streptactin resin during purification. The protein was subjected to a cycle of MT binding and release by ATP to select for active motors. Briefly, motors were bound to an excess of stabilized MTs in BRB80 buffer with 10 μM taxol. MTs were pelleted at 60,000 x g for 10 min at room temp. Bound motors were released by resuspension of the MT pellet in BRB80 with 10 μM taxol and 10 mM ATP. MTs were pelleted again as before and the eluted motors were frozen in LiN_2_ after the addition of 20% sucrose and 1 mg/ml BSA as cryoprotectants.

The DDB complex was prepared by adding recombinant labeled strepII-SNAPf tagged BiCD2 (N-terminal construct encompassing amino acids 25-400) to high-speed porcine brain lysates as previously described (McKenney et al., 2014). The DDB complexes were fluorescently labeled with excess SNAP-Cell TMR-Star dye (NEB) during purification as described (McKenney et al., 2014) and aliquots of eluted DDB were flash frozen in LiN_2_ and stored at −80^o^C. We note that freezing the complex leads to an apparently larger percentage of diffusive complexes in our assays (~15% for unfrozen versus ~30% for frozen).

### Microscopy experiments and quantification

Glass chambers were prepared by acid washing as previously described (Tanenbaum et al., 2013). Polymerized microtubules were flowed into streptavidin adsorbed flow chambers and allowed to adhere for 5-10 min. After washing the excess unbound microtubules using assay buffer (30 mM Hepes pH 7.4, 50 mM K-acetate, 2 mM Mg-acetate, 1 mM EGTA, 10% glycerol, 0.1 mg/mL biotin-BSA, 0.2 mg/mL K-casein, 0.5% Pluronic F127, and an oxygen scavenging system (Aitken et al., 2008)), a motility mixture containing labeled DDB complex or dynactin (p150 and p135) or recombinant GST-hDyn was then flowed as described earlier (McKenney et al., 2014). Images were acquired using Micromanager software (Edelstein et al., 2010) controlled Nikon TE microscope (1.49 N.A., 100x objective) equipped with a TIRF illuminator and Andor iXon CCD EM camera. In the case of GST-hDyn, or DDB complex, 2 mM ATP was included in the buffer. Velocities were calculated from kymographs generated in ImageJ. For fluorescent intensity values we used maximum intensity projections of time series to quantify GST-hDyn due to its transient binding to the MT. For p150 and p135, raw images, were quantified due to these proteins longer bound lifetime on the MT. Standard deviation maps (Cai et al., 2010; Cai et al., 2009) were generated using the image stack Z-projection function in ImageJ. For figure preparation, microscopy images were background subtracted using the ‘subtract background’ function in ImageJ with a rolling ball radius of 50 pixels. Image contrast was linearly adjusted using ImageJ. Because the DDB molecules often run the entire microtubule lengths, we did not analyze the run-lengths of DDB motility.

The authors declare no competing financial interests.

## SUPPLEMENTAL INFORMATION

**Figure S1.**
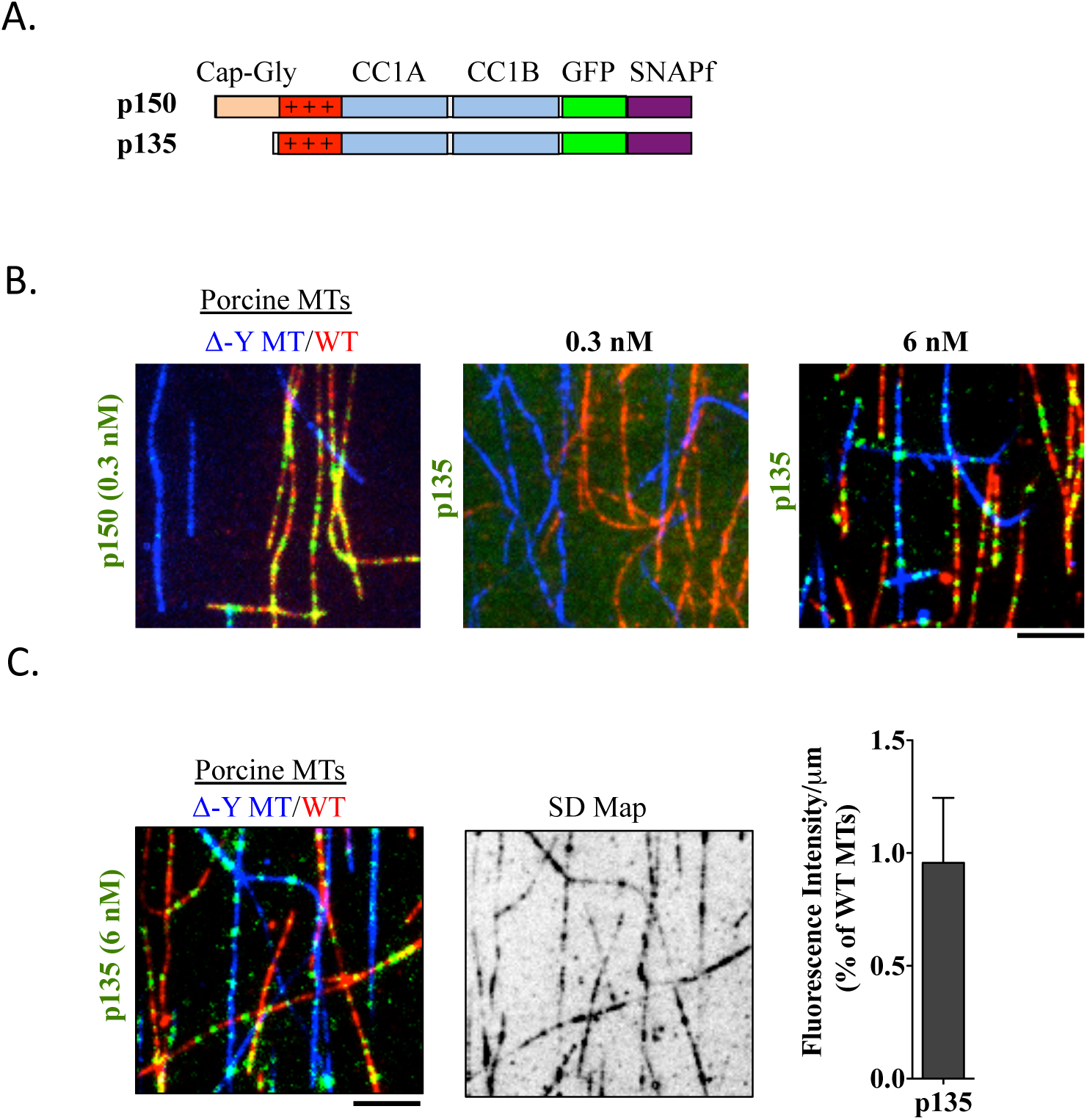
Role of the p150 CAP-Gly domain in recognition of α-tubulin tyrosination. **(A)** Schematic of the p150 constructs used. The p135 isoform lacks the CAP-Gly domain, but retains the adjacent basic domain (red) and coiled-coil domains (blue). (B) TIRF image of either p150 (0.3 nM), or p135 (0.3 or 6 nM) bound to either WT (red), or CPA-treated porcine MTs (blue). Note: no binding of p135 is observed at comparable concentrations to p150 indicating the lower binding affinity of this construct. Proteins were visualized by fluorescence from the GFP tag fused to each construct. (C) Still images from a TIRF time-series of p135 binding to either WT (red), or CPA-treated porcine MTs (blue). Right, standard deviation map of p135 binding shows no preference for either type of MT, and quantification of mean fluorescence intensity (arbitrary units) per μm MT for p135 bound to CPA-treated MTs relative to WT MTs in the same chamber (n ≥ 40 MTs quantified for each condition from two replicate experiments). Scale bars are 5 μm.

**Figure S2.**
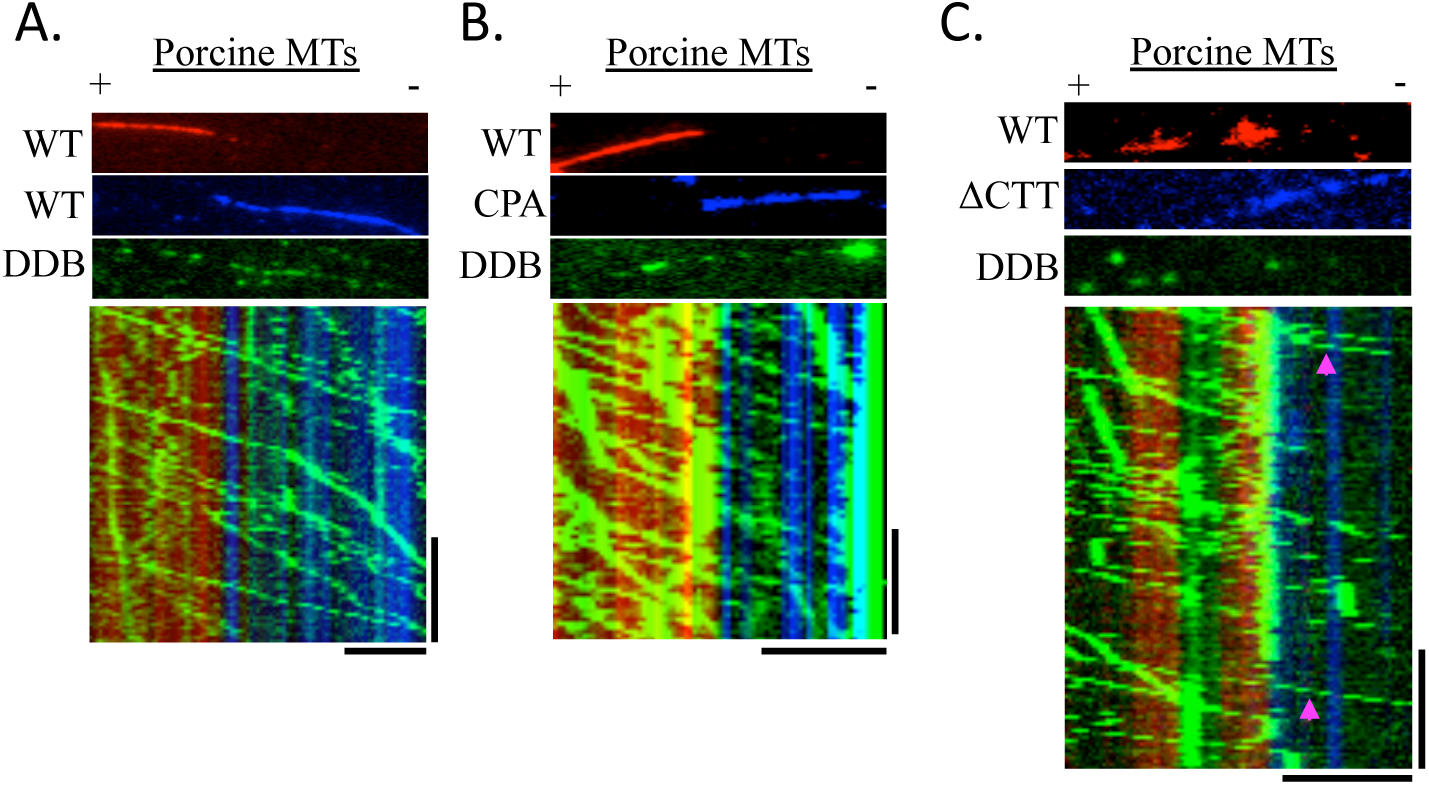
Continuous DDB movement does not require tyrosination or tubulin CTTs. **(A-B)** TIRF images and kymographs of annealed WT MTs **(A)**, or WT and CPA-treated porcine MTs **(B)**. Note TMR-labeled DDB complexes (green) in **(A)** traverse the annealed boundary without pausing (93.5%, n=356 processive complexes, 23 MT-MT junctions, 2 replicate experiments), suggesting that the annealing process does not create a barrier to DDB motility. (B) The majority of DDB complexes (77.6%, n=960 processive complexes, 47 MT-MT junctions, 2 independent experiments) move processively across the boundary between WT and CPA-MTs. **(C)** TIRF image and kymograph from a chimeric MT composed of WT (red) and subtilisin-treated porcine MTs (blue). Note some DDB complexes (pink arrows) traverse the boundary and continue moving along the subtilisin-treated section of the MT (41.1%, n=709 processive complexes, 56 MT-MT junctions, 3 independent experiments). All scale bars are 5 μm and 25 sec.

**Video S1. Comparison of DDB motility on recombinant MTs.** Movie shows movement of fluorescent TMR-labeled DDB molecules (green) on either WT recombinant yeast MTs (red), or on MTs lacking the α-tubulin (Del-A, blue MTs), β-tubulin (Del-B, blue), or tail-swapped MTs (Tail Swap, blue) placed in the same chamber.

**Video S2. Comparison of DDB motility on tyrosinated and detyrosinated MTs.** Movie shows movement of fluorescent TMR-labeled DDB molecules (green) on either fully tyrosinated (red), or detyrosinated (blue) recombinant yeast MTs. Scale bar is 5 μm.

**Video S3. DDB motility on chimeric MTs.** Movie shows movement of fluorescent TMR-labeled DDB molecules (green) on a single chimeric MT composed of tyrosinated (red) and detyrosinated (blue) MT sections. Note that processive DDB molecules traverse the junction between tyrosinated and detyrosinated MT sections and continue processive movement, while diffusive complexes do not cross the boundary. Scale bar is 5 μm.

